# Priority Effects Can Shape Strain-Level Community Assembly and Function in a Bioluminescent Symbiosis

**DOI:** 10.1101/2025.05.01.651688

**Authors:** Emma D. Román, Andrea N. Nebhut, Tadashi Fukami, Alison L. Gould

## Abstract

Symbiotic microbial communities often appear highly variable in their composition and function in ways that environmental factors alone cannot explain. One potential reason for this variability is priority effects, where historical contingency in arrival order affects how symbionts assemble into communities. Focusing on the luminous bacterium (*Photobacterium mandapamensis*) in the light organ of the sea urchin cardinalfish (*Siphamia tubifer*), we studied how priority effects might influence bacterial symbiont assembly. In *in vitro* experiments that used three *P. mandapamensis* strains isolated from the same individual fish, we found that whichever strain arrived first dominated over the other two strains, indicating strong priority effects. We also found that the strains differed in growth and luminosity, and that the bioluminescence of the three-strain community could not be predicted from individual strain performances. These findings suggest that priority effects can be a major process shaping the composition and function of these symbiotic microbial communities.

## Introduction

The assembly of symbiotic microbial communities is complex, with interactions among the hosts, the symbionts, and the environment all playing a role [1–4]. One factor that can drive this assembly is the arrival order and timing of different taxa [5]. Taxa that arrive early can outcompete later-arriving taxa by occupying available niches and modifying the local environment to facilitate or inhibit the growth of subsequent taxa. A limited number of recent studies indicate that these historical influences, known as priority effects, can play a major role in shaping symbiotic communities made of different strains [6–15].These studies show that priority effects can operate not just among species, but also among strains. However, priority effects remain relatively understudied as a factor affecting symbiotic communities.

We examine priority effects in a luminous bacterial symbiont, *Photobacterium mandapamensis*. Their host, *Siphamia tubifer*, is an Indo-Pacific cardinalfish that resides among the spines of sea [16, 17]. Each fish has a specialized, gut-associated light organ that houses *P. mandapamensis* [18, 19]. The bacterially produced light is used when fish leave the urchin at night to forage [18, 19], presumably for camouflage by counterillumination. This symbiosis begins about one week after larval development, and *P. mandapamensis* is the only bacterium known to colonize the light organ of *S. tubifer*, forming a specific binary association [18, 20]. *Photobacterium mandapamensis* is considered a subspecies to *Photobacterium leiognathi* which further highlights the specificity of the *S. tubifer–P. mandapamensis* symbiosis [21].

Despite this binary relationship, *S. tubifer* light organs contain an average of five to six *P. mandapamensis* strains, with minimal overlap between individual hosts (Gould et al., 2023). Each *P. mandapamensis* strain is highly similar with an average nucleotide identity of greater than 97%, indicating that strains might have overlapping niches [22]. Despite this, different bacterial genotypes (i.e. strains) can exhibit phenotypic differences in traits such as metabolic capacity, competitive strategies, and host specificity [23–26]. Such variations may influence symbiont community structure and function, yet the role of naturally co-occurring, closely related strains within symbioses remains underexplored [24, 25, 27, 28]. This juxtaposition of host specificity and strain-level diversity raises compelling questions about how these communities assemble, and whether variation in strain composition influences the functional outcome of the symbiosis.

The natural simplicity of the *S. tubifer–P. mandapamensis* partnership positions it as a unique model for investigating whether high strain diversity has functional consequences for the host [29]. Building on previous findings that individual hosts harbor distinct sets of closely related strains [20], we asked whether priority effects contribute this variation, if strains differ from each other in ecologically meaningful ways, and does community composition impact function. To address these questions, we conducted *in vitro* assembly experiments using three *P. mandapamensis* strains isolated from the same individual fish [20], and we quantified growth rate and bioluminescence in mono- and mixed cultures.

## Methods

### Bacterial Strains and Fluorescent Tagging

This study used three *P. mandapamensis* strains, Ph.E, Ph.O, and Ph.P, isolated from the light organ of the same S. *tubifer* individual collected from Verde Island in the Philippines [20]. These bacterial strains were identified through PCR fingerprinting in a previous study [20]. To enable tracking in co-culture experiments, Ph.E and Ph.O were engineered to constitutively express the fluorescent proteins mVenus and mRuby, respectively, by San Diego State University’s Research Foundation (SDSURF) using established methods [30, 31]. Fluorescent tagging was confirmed not to impact growth rates and allowed for strain tracking during competition assays (Figure S1).

### Strain Phenotype Assays

#### Growth Rate

Growth rates were measured using a BioTek Synergy H1 Hybrid Multi-Mode Microplate Reader and BioTek Gen5 Microplate Reader and Imager software (v3.10). Wild-type strains were revived from -80°C glycerol stocks, streaked onto agar plates containing Lennox broth in 70% seawater (hereby LSW-70), and incubated at 26°C for 24 hours. Single colonies of each strain were inoculated into 3 mL of LSW-70 liquid media and grown overnight. Cultures were then standardized to an OD600 of 0.7. Afterwards, cultures were diluted 1:100 and added to a 96-well plate. 100 μL of uninoculated media was used as blanks. Absorbance readings at 600 nm were then taken every 5 minutes for 24 hours at room temperature with the lid on to reduce evaporation. Growth curves were analyzed in R (v4.2.1) using the growthcurver package [32], and growth rates (r) were compared using an ANOVA with Tukey-adjusted post hoc comparisons of estimated marginal means.

#### Luminosity

Luminosity was quantified for each strain individually and across multiple community compositions. To compare individual strains, liquid cultures of wild-type strains were grown overnight in LSW-70 media at 26°C with shaking (120 rpm) in a New Brunswick Innova 43/43 R Console Incubator Shaker. Cultures were standardized to an OD600 of 0.9, and 100 μL of each overnight culture was then used to inoculate 10 tubes with 3 mL of LSW-70 broth (n = 10 for each strain). After 15 hours, cultures were adjusted to an OD600 of 0.7 to ensure comparable cell densities before luminescence measurements.

To compare the luminosity of communities, eight individual isolates of each focal strain were picked and grown in liquid cultures overnight at 26°C. Two isolates for each strain were grown in 25 mL of liquid media and used for both individual strain readings and to create mixed cultures. The remaining six isolates were grown in 3 mL and used solely for individual strain measurements. After 15 hours, cultures were adjusted to an OD600 of 0.7 using a Thermo Fisher Varioskan ALF Multimode Microplate Reader with accompanying SkanIt software. Mixed community compositions were generated by inoculating each strain at either a high or low initial density to mirror the conditions used in the competition assays. Communities were assembled by adding 1000 μL of a strain for high-concentration treatments or 10 μL for low-concentration treatments and then mixed. All possible combinations of the three strains were used.

Luminescence was measured immediately (within 30 miuntes) after the preparation of single- and multi-strain replicates using a GloMax 20/20 Luminometer (Promega). For each sample, 1 mL aliquots were shaken for 5 seconds before each reading, and luminosity was reported in relative light units (RLUs). All measurements were standardized to blank measurements, which contained 1 mL of LSW-70 broth. Data were analyzed in R (v4.2.1), with strain comparisons performed using ANOVA and post hoc estimated marginal means with Tukey HSD correction.

### Mix-culture Experiments

#### Sample Preparation

Overnight cultures of the fluorescently labeled Ph.E, Ph.O, and wild-type Ph.P strains were grown as previously described and standardized to an OD600 of 0.7. To investigate priority effects, altered starting concentrations were used to simulate differences in arrival order. High (∼10^4^ CFU/mL) and low (∼10^2^ CFU/mL) initial concentrations were prepared through serial 1:10 dilutions, serving as proxies for “early” and “late” arrivals, respectively. For altered timing experiments, strains at high concentration (∼10^4^ CFU/mL) were introduced sequentially to simulate staggered arrival. The high concentration from the altered starting concentration experiments was used for altered timing experiments to optimize the efficiency of flow cytometry data collection.

#### Pairwise Experiments

Pairwise combinations of strains at high and low starting concentrations were used to determine the effect of arrival order on strain abundances. Experiments were conducted in clear, flat-bottom 96-well plates. Each well received 80 μL of sterile LSW-70 media, with strain combinations added as 10 μL aliquots (Table S1). Controls included wells containing only one strain at a 1:10 dilution or blank wells with 100 μL of sterile LSW-70 media. Absorbance was recorded every 5 minutes for 24 hours using a BioTek Gen5 Microplate Reader. The lid of the 96-well plate was kept on throughout the experiment to reduce evaporation. At 0, 10, and 24 hours, wells were destructively sampled in triplicate and diluted as follows: 1:10 at 0 hours and 1:10^6^ at 10 and 24 hours. Diluted samples were plated on LSW agar and incubated at 26°C for 24 hours.

#### Mixed Community Experiments

To investigate priority effects in a community with all three strains present, combinations of high and low starting densities were tested in clear, flat-bottom 96-well plates (Table S2). Wells were prepared with 70 μL of sterile LSW-70 media, and 10 μL of high (∼10^4^ CFU/mL) or low (∼10^2^ CFU/mL) inoculum was added for each strain. Controls included wells containing only one strain or blank wells with 100 μL of sterile LSW-70 media. The absorbance of the inoculated 96-well plates was then measured with a BioTek Gen5 Microplate Reader at room temperature over 24 hours. The lid of the 96-well plate was kept on throughout the experiment to reduce evaporation. Wells were destructively sampled in triplicate at 0, 10, and 24 hours, following the same dilution and plating protocols as described above.

Colonies from the pairwise and three-way experiments were enumerated using a Dino-Lite fluorescence handheld microscope. Strain abundance was measured by colony counts. Strains with the low starting concentration produced too few colonies to calculate CFU/mL due to plating limitations. When zero colonies were counted, counts were reassigned a value of 1 to reflect very low abundance rather than true absence. Based on the design of the experiment, differences between treatments were large enough that meaningful patterns could still be interpreted.

#### Flow Cytometry Experiments

Flow cytometry was employed to achieve higher-resolution analysis of strain abundances, focusing on either altered strain concentrations or staggered strain arrival orders. Experimental setups followed the three-way protocol, using fluorescently labeled Ph.E and Ph.O strains (Tables S3 & S4). Two 96-well plates were used to allow for three replicates per time point, and both 96-well plates were grown in a New Brunswick Innova 43/43 R Console Incubator Shaker at 120 rpm and 26 °C. Optical density readings were taken every hour for 12 hours and once at 24 hours using a BioTek Synergy H1 Hybrid Multi-Mode Microplate Reader. Samples were analyzed at 4, 6, 8, 10, 12, and 24 hours using an Attune NxT Flow Cytometer. Instrument settings were optimized for mRuby and mVenus fluorescence, and gating strategies were applied to isolate single-cell events. Blank wells containing sterile LSW-70 media and control wells containing each strain grown separately were analyzed to ensure minimal background fluorescence. Samples were diluted 1:10 in PBS before analysis, except for 4-hour samples, which were run undiluted. A 10% bleach solution was used between samples to minimize contamination. Each sample was passed through the flow cytometer at a rate of 12.5 μL/sec until 100,000 events were collected. Afterwards, the abundance of non-fluorescent, mRuby, and mVenus populations was recorded.

Flow cytometry data were prepared using the FlowJo software (v10.10, BD Biosciences). All experiments received the same gating parameters, and counts for each strain were then exported for analysis. Bacterial abundance (counts/mL) was calculated using the same approach as CFU/mL determination. One biological replicate from the 10- and 12-hour timepoints of the altered timing experiment were excluded due to a technical error that resulted in a missing signal for one or more strains.

To assess whether differences in initial abundance or arrival order influenced community composition, bacterial counts over time were analyzed using linear models in R (v4.2.1). Furthermore, the relative abundance of each strain within each community was analyzed at the 10-hour timepoint to be consistent with the plating data presented in the Supplementary Information. The 10-hour data was chosen to minimize the influence of dead cells, which are more prevalent at later timepoints. Technical replicates were averaged, and log-transformed counts/mL were modeled using linear models that included strain identity and a three-way interaction among the initial concentrations (or arrival order) of Ph.E, Ph.O, and Ph.P. Differences in relative abundance were evaluated using ANOVA, followed by post hoc pairwise comparisons of estimated marginal means with Tukey’s HSD correction.

## Results

### Strains Differ in Growth Rate and Luminosity

The strains differed significantly in growth rate (r) and luminosity (relative luminosity units, RLU) (Figure 1). Ph.E had an average growth rate of 0.83 and grew significantly faster than Ph.O (r = 0.47, p < 0.001) and Ph.P (r = 0.52, p < 0.001). Ph.P and Ph.O did not differ significantly in growth rate (p = 0.59). Ph.E was the fastest-growing strain on average, but it also exhibited the most variability, with a minimum of 0.59 and a maximum of 0.99. The slowest individual growth rate observed across all strains was 0.40 for Ph.O.

**Figure 1:**
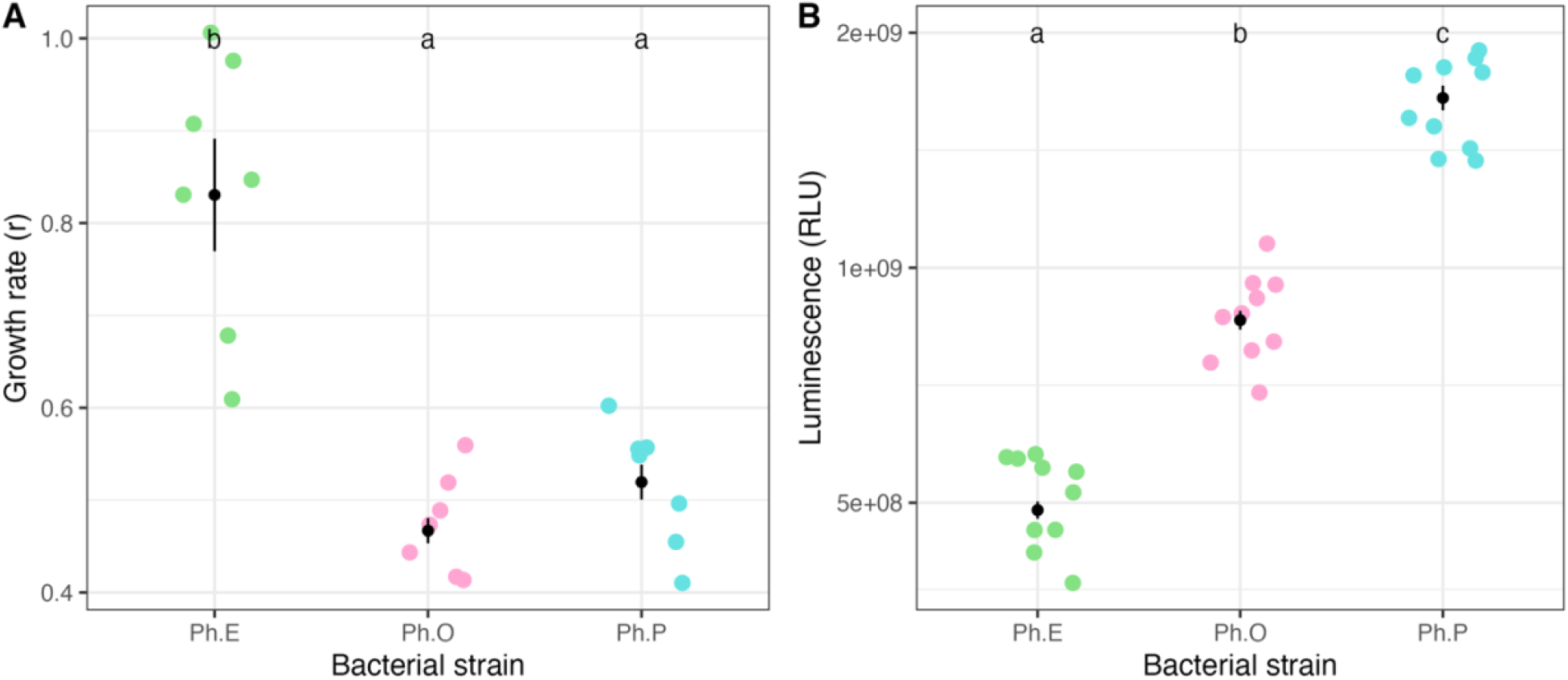
Growth rate (r, *A*) and luminescence (relative luminosity units, RLU, *B*) of *Photobacterium mandapamensis* strains Ph.E (green), Ph.O (pink), and Ph.P (blue). Individual data points (n = 10) are shown with means ± standard error indicated by black bars. Strains sharing the same letter are not significantly different from one another (p > 0.05). Ph.E exhibited the highest growth rate, while Ph.P showed the highest luminosity.

In contrast to growth rate, Ph.E was the darkest strain with an average luminosity of 4.9 × 10^8^ RLUs and the lowest observed luminosity of 4.4 × 10^8^ RLUs. Ph.P was the brightest strain, with an average luminosity of 1.7 × 10^9^ RLUs, followed by Ph.O (8.6 × 10^8^ RLUs). Unlike growth rate, all three strains differed significantly in luminosity (Ph.O vs. Ph.E p < 0.001, Ph.P vs. Ph.E p < 0.001, and Ph.O vs. Ph.P p < 0.001). Ph.P exhibited the greatest variability in light production, ranging from 1.5 × 10^9^ RLUs to 1.9 × 10^9^ RLUs.

### Strain Mixing Influences Luminosity

Luminosity varied significantly across communities with different strain compositions (Figure 2, Table S5). We recreated the naturally occurring community composition described in Gould et al. (2023), using 81% Ph.E, 5% Ph.O, and 14% Ph.P, and compared it with experimental communities simulating differences in arrival order with differences in the starting concentration of each strain.

**Figure 2:**
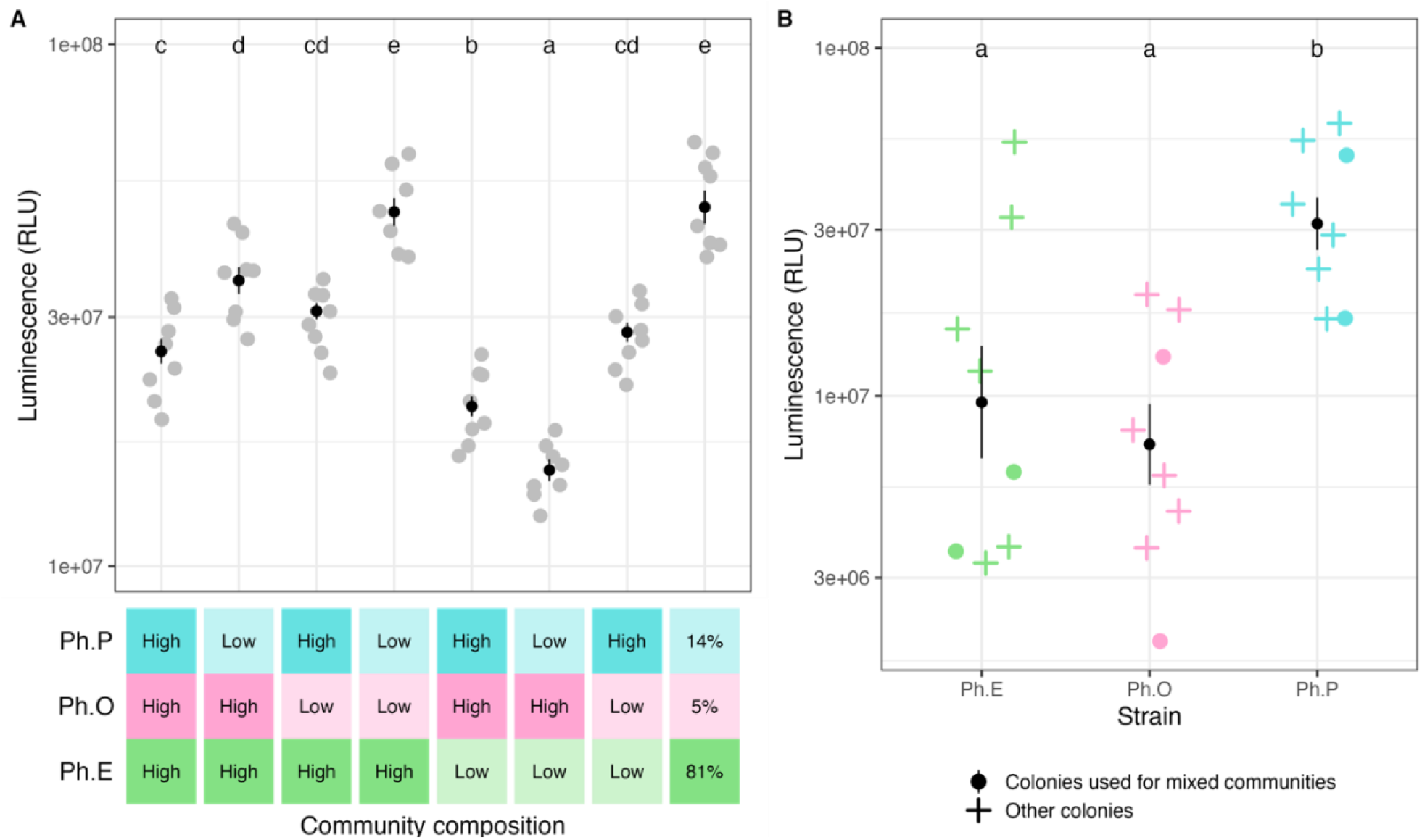
Luminescence (relative luminosity units, RLU) of *Photobacterium mandapamensis* strains Ph.E (green), Ph.O (pink), and Ph.P (blue) grown either in mixed communities (A, gray points) or individually (B). Individual colonies of each strain used to create mixed communities (circles) are differentiated from colonies just measured in isolation (plus). Community compositions are listed on the x-axis using high or low concentrations, and the naturally occurring community was recreated using 81% Ph.E, 5% Ph.O, and 14% Ph.P (Gould *et al*. 2023). Individual data points (n = 10 and n = 8, respectively) are shown with means ± standard error (black bars). Strains sharing the same letter are not significantly different (p>0.05). When Ph.E is grown at high concentration in a mixed community, the community is bright.

As in Figure 1, Ph.P was significantly more luminous than both Ph.E (p=0.02) and Ph.O (p=0.004) when strains were measured in isolation (Figure 2B). The recreated natural community (81% Ph.E, 5% Ph.O, and 14% Ph.P) was the most luminous overall, with an average of 5.0 × 10^7^ RLUs, closely followed by the Ph.E-dominated experimental community at 4.8 × 10^7^ RLUs. Both were significantly more luminous than all other community compositions (Figure 2A, Table S5). Ph.E was the most abundant strain in both communities.

Communities with low concentrations of Ph.E were consistently dimmer, with the Low–High– Low treatment being the darkest (average = 1.6 × 10^7^ RLUs). Although Ph.P was the brightest strain when grown in isolation, communities where it was most abundant were not especially bright (Figure 2).

### Priority Effects Influence Community Assembly

After growing in mixed culture, the final community composition closely reflected the initial conditions (Figure 3; Figures S3, S4, & S5). In both pairwise and three-strain communities, priority effects shaped final community composition (Figures S3 & S4). After 10 hours, the strains introduced with a high initial concentration were consistently most abundant (Figures S3 & S4). While strains introduced at low concentration remained detectable, too few colonies were counted to reliably estimate CFU/mL (Figures S3 & S4).

**Figure 3:**
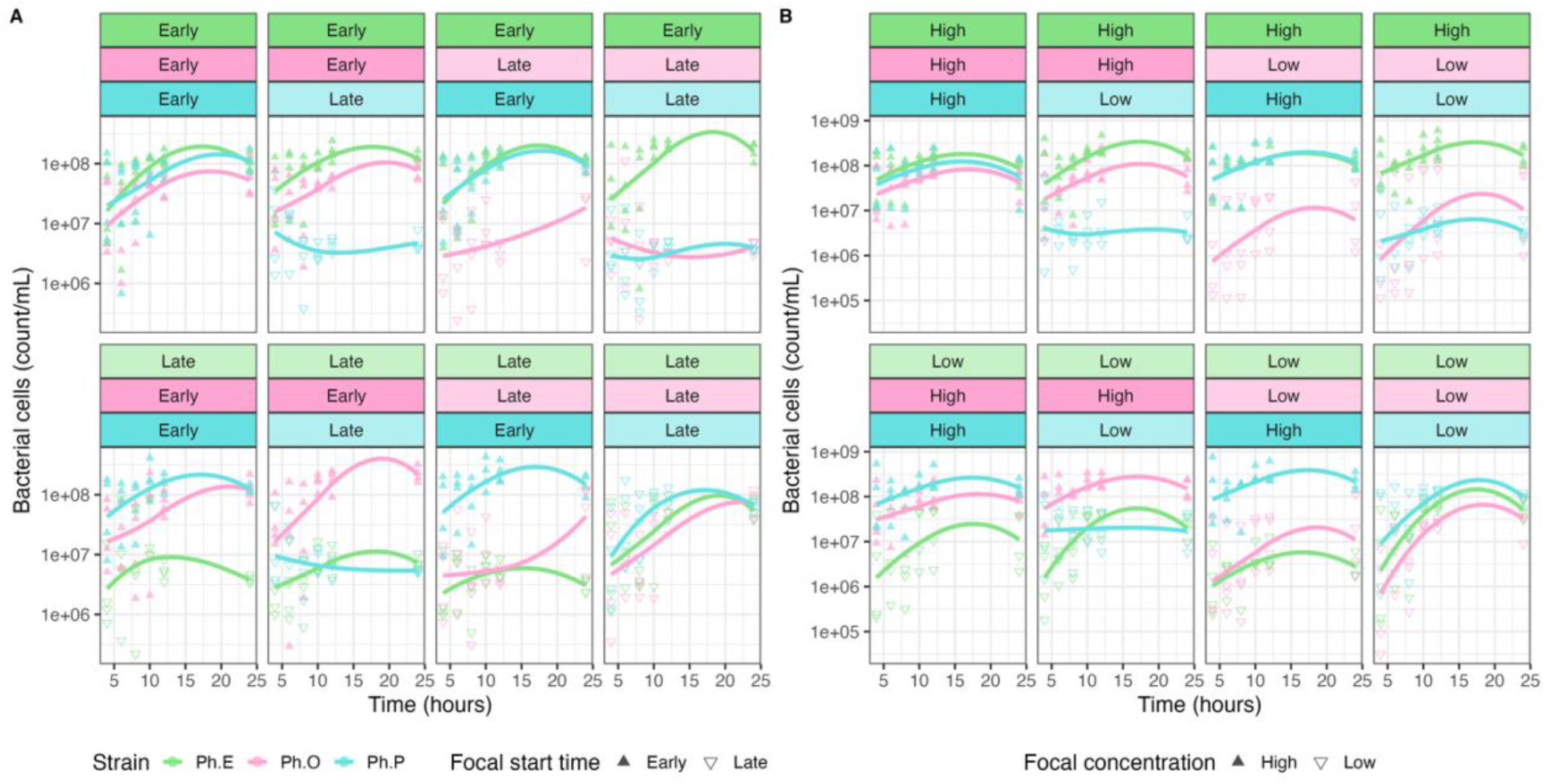
Abundance of *Photobacterium mandapamensis* strains Ph.E (green), Ph.O (pink), and Ph.P (blue) when grown in mixed communities over 24 hours under different initial conditions. Strain abundance was measured using flow cytometry. Priority effects were simulated by staggering arrival order (A) or altering initial strain concentrations as a proxy for arrival order (B). The treatment conditions are indicated above each panel. Upward-pointing triangles denote strains that arrived first or were introduced at a higher concentration; downward-pointing triangles indicate strains that arrived later or at a lower concentration. Each point represents the average abundance across biological replicates (n = 3 for the all-late treatment at 10 and 12 hours, n = 4 for all other treatments and timepoints), with a loess-smoothed best-fit curve.

When measured using flow cytometry, priority effects remained strong, and community structure remained relatively constant over time, regardless of whether strain arrival order or initial concentration was manipulated (Figure 3). Due to increased cell death creating more noise at later time points, we highlight the 10-hour results as they provide a clearer snapshot of community composition. However, community composition after 24 hours showed consistent trends (Figure S4). Across treatments, no single strain consistently outcompeted the others, and there were no discernible differences between experiments manipulating arrival order versus initial concentration (Figure 4).

**Figure 4:**
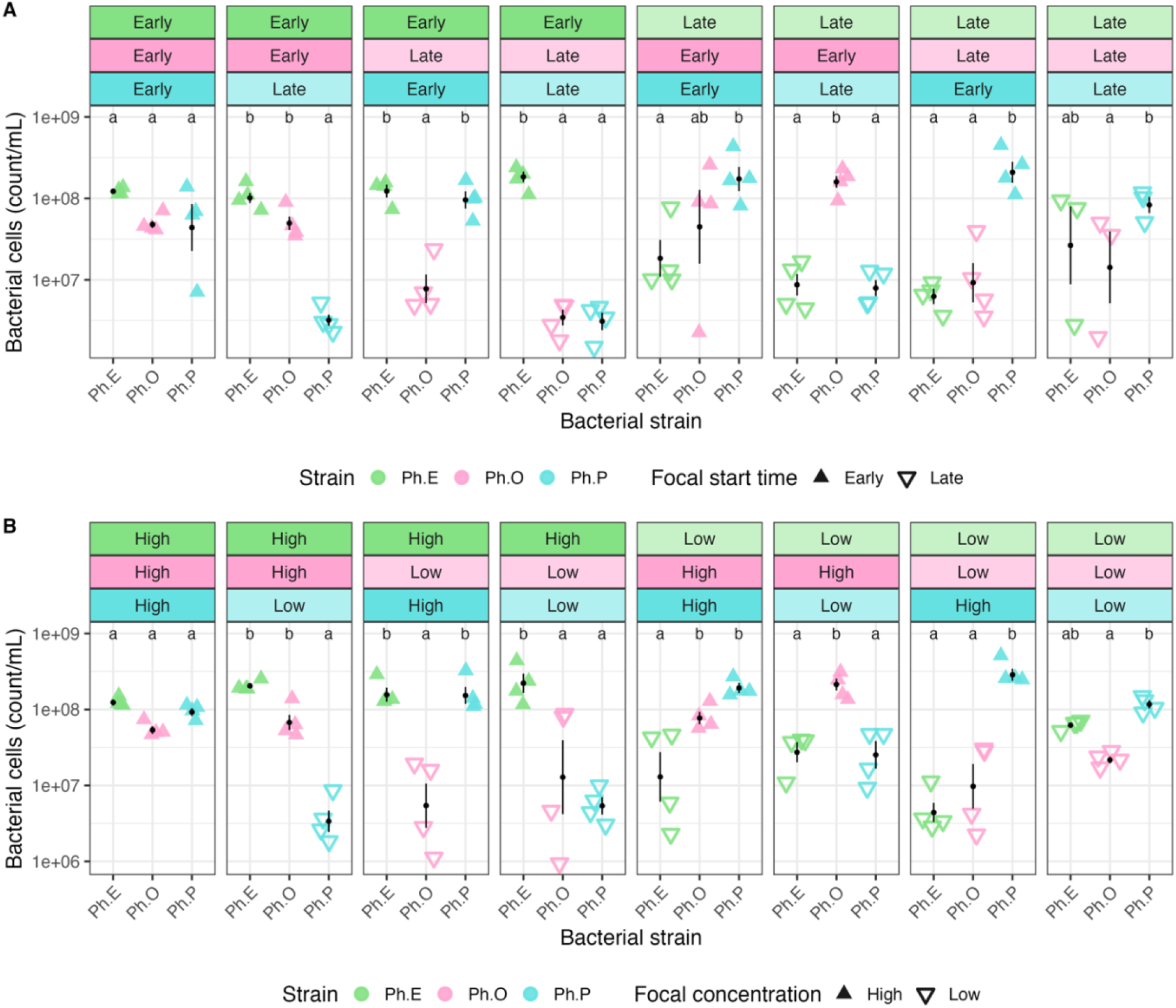
Abundance of *Photobacterium mandapamensis* strains Ph.E (green), Ph.O (pink), and Ph.P (blue) when grown in mixed communities after 10 hours under different initial conditions. Strain abundance was measured using flow cytometry. Priority effects were simulated by staggering arrival order (A) or altering initial strain concentrations as a proxy for arrival order (B). The treatment conditions are indicated above each panel. Upward-pointing triangles denote strains that arrived first or were introduced at a higher concentration; downward-pointing triangles indicate strains that arrived later or at a lower concentration. Each point represents the average abundance across biological replicates (n = 4, n = 3 for the all-late treatment), with black points representing means ± standard error indicated by black bars. Strains sharing the same letter are not significantly different from one another (p>0.05).

Ph.O exhibited the greatest variability in abundance across treatments (Figure 4). For instance, Ph.O was less abundant than Ph.P when all strains were introduced late (*p* = 0.04) or at low concentrations (*p* = 0.01). These differences were no longer apparent at the 24-hour time point (Figure S4). Additionally, Ph.O was not significantly more abundant than Ph.E when Ph.E was introduced late. However, under the equivalent conditions when concentration was manipulated, Ph.O was significantly more abundant than Ph.E (*p* = 0.01). Though these differences were not statistically significant, the abundance of Ph.O was also consistently equal to or higher than Ph.E when both strains were introduced late or at low concentrations (Figure 4). This trend was also reflected in the 24-hour data (Figure S4). Ph.P exhibited the most variability when all strains were introduced early, but remained highly consistent across all other treatments.

## Discussion

Our results suggest that naturally co-occurring strains of *P. mandapamensis* vary in growth and luminosity (Figure 1) and that arrival order exerts a strong influence on community assembly (Figure 4). Previous work has shown that individual hosts harbor unique strain communities with limited sharing across individuals [20]. Our findings offer a possible explanation for this pattern, i.e., that differences in arrival history underlie the high strain variation between hosts. For example, the dominance of strain Ph.E over Ph.O and Ph.P observed in one fish [20] may represent the historical contingency of Ph.E happening to have arrived early. In this system, symbiont acquisition begins at least seven days after larvae become pelagic, during which time larvae are likely exposed to a diverse environmental pool of potential symbionts before they settle onto urchin habitats three weeks later [18]. Therefore, it is possible that this fish could have had a different dominance pattern if the assembly history were different. Arrival history can be particularly variable in horizontally transmitted symbioses, such as the *S. tubifer–P. mandapamensis* association, where symbionts are acquired entirely from the environment, increasing the likelihood that arrival timing shapes community assembly [6, 25, 33].

Across all experimental conditions, our results were consistent (Figures 3 & 4; Figures S3, S4, & S5). Strains that arrived early or had the highest initial abundance consistently became the most abundant (Figure 4). The strong priority effects found in our study may be due to high niche overlap, which can intensify priority effects among closely related taxa [34]. In symbiotic systems, the number of co-inhabiting conspecific strains seems to have ecological constraints. For example, humans typically harbor 1–4 strains per species in the gut microbiome [35, 36], Hawaiian bobtail squid host 1-6 strains of *Vibrio fischeri* within their light organ [37], and *S. tubifer* hosts on average 5–6 *P. mandapamensis* strains [20]. The high inter-host variation of the 5–6 strains might allude to a host’s capacity to acquire multiple strains, with priority effects dictating which strains dominate. A similar mechanism appears to happen in the squash bug– *Caballeronia* symbiosis, where high strain diversity across hosts has been linked to strong priority effects of horizontally acquired symbionts [38].

By independently measuring growth rate and bioluminescence, we found variation across strains, including a potential trade-off between these traits (Figure 1). This was most apparent in strain Ph.P, which had the lowest growth rate and highest luminescence (Figure 1). Though we also find that luminosities of each strain were variable (Figures 1 & 2B), our results demonstrate clear differences in luminescence among strains. These findings build on previous work showing similar dynamics in other strain combinations [20] and suggest that conspecific strains can differ in functional traits.

Furthermore, bioluminescence of strains in isolation did not always predict community-level light output (Figure 2). For example, communities where Ph.E was most abundant, including the recreated natural community, were the brightest (Figure 2A) despite its lower luminescence when measured alone. Since luminescence was measured immediately after mixing strains, these results suggest that strain interactions affecting light production can occur rapidly. Although the reason for this result remains unclear, Ph.E may produce more light in the presence of Ph.O and Ph.P, potentially through differences in metabolite chemistries. Such dynamics may reflect a form of metabolic division of labor, in which strains perform complementary roles that reduce individual metabolic burdens while sustaining overall community output [39]. Future experiments measuring luminescence of mixed communities and individual strains over time will help clarify the timing, stability, and mechanisms underlaying light production.

*Siphamia tubifer* is believed to rely on symbiont-produced bioluminescence while foraging at night, making light production essential to host fitness [18]. As shown in this study and others, however, there is likely an energetic cost for bacteria to create light (Figure 1) [40, 41]. Strain diversity introduced by priority effects may increase functional redundancy which in turn could help maintain host fitness and a stable symbiotic partnership [39, 42, 43]. Measuring the luminosity of other natural strain communities within this system could reveal whether community-level light production is consistent across hosts. If so, it would reinforce the idea that microbial interactions influence host fitness [4].

To better understand priority effects in this system, *in vivo* experiments that build on the *in vitro* study presented here are necessary. In *S. tubifer*, the light organ consists of many chambers housing *P. mandapamensis* [17, 18]. A similar structure exists in the Hawaiian bobtail squid [44]. In this system, the light organ has six crypts, which strains of *V. fischeri* rarely coinfect, suggesting spatial partitioning influences strain coexistence [24, 45]. Similar partitioning may occur in the *S. tubifer* light organ, where priority effects could unfold independently across multiple chambers contributing to the high strain-level diversity observed across hosts [20].

In conclusion, this study suggests that priority effects influence community composition and function among naturally co-occurring strains of *P. mandapamensis* (Figures 3 & 4). We found significant phenotypic variation between strains in growth and bioluminescence (Figure 1), yet community composition was a better predictor of total light output than the performance of the dominant strain alone (Figure 2). This suggests that priority effects shape symbiont communities which influences bioluminescent function, a key feature of this symbiosis. Our results lay a foundation for future *in vivo* investigations into how compartmentalization, host filtering, and strain interactions together shape symbiont communities alongside priority effects.

## Acknowledgements

We thank Amaury Payelleville and Jean Villa for suggestions on experimental design, Tatjana Schlechtweg for assistance with the flow cytometer, Amir Yarmahmoodi for access to FlowJo, Joy Kumagai for suggestions on statistical analysis, and Lilly Hammer for help collecting luminosity data.

## Funding

Funding was provided by the National Institute of Health (DP5-OD026405).

## Author Contributions

ER, AG, and TF designed the study. ER conducted experiments, analyzed data with AN, and wrote the first draft of the manuscript. AN performed bioinformatic analysis and prepared figures. AG secured funding for this work. All authors contributed to revisions of the manuscript.

## Supplementary

**Figure S1:**
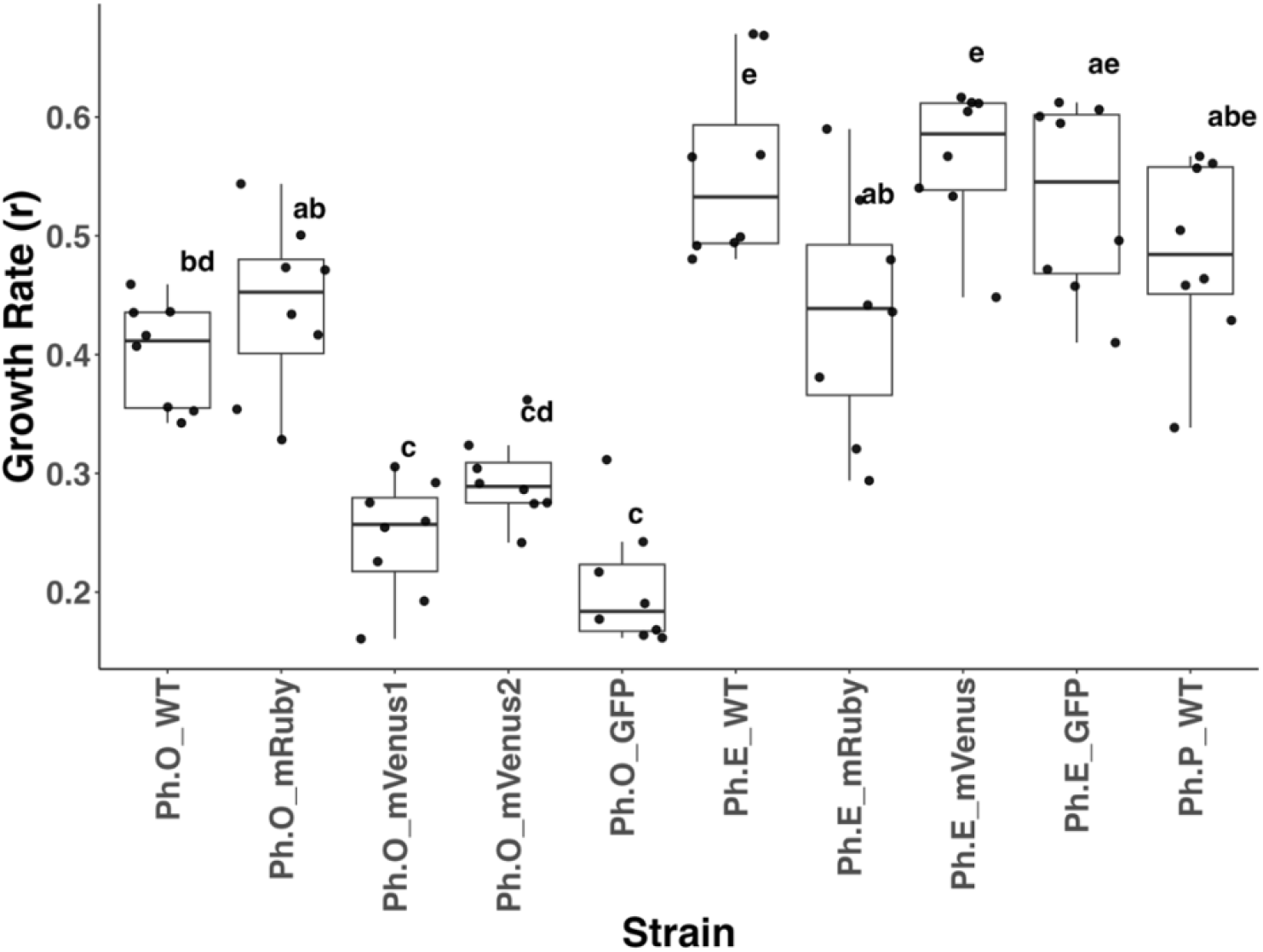
A box plot showing the growth rates for strains Ph.P, Ph.O, and Ph.E with various fluorescent tags. Strains sharing the same letter are not significantly different from one another (p>0.05). Ph.P did not have any fluorescent tags, so the wild type was chosen. For Ph.O, the mRuby tag was chosen, and for Ph.E the mVenus tag was chosen.

**Figure S2:**
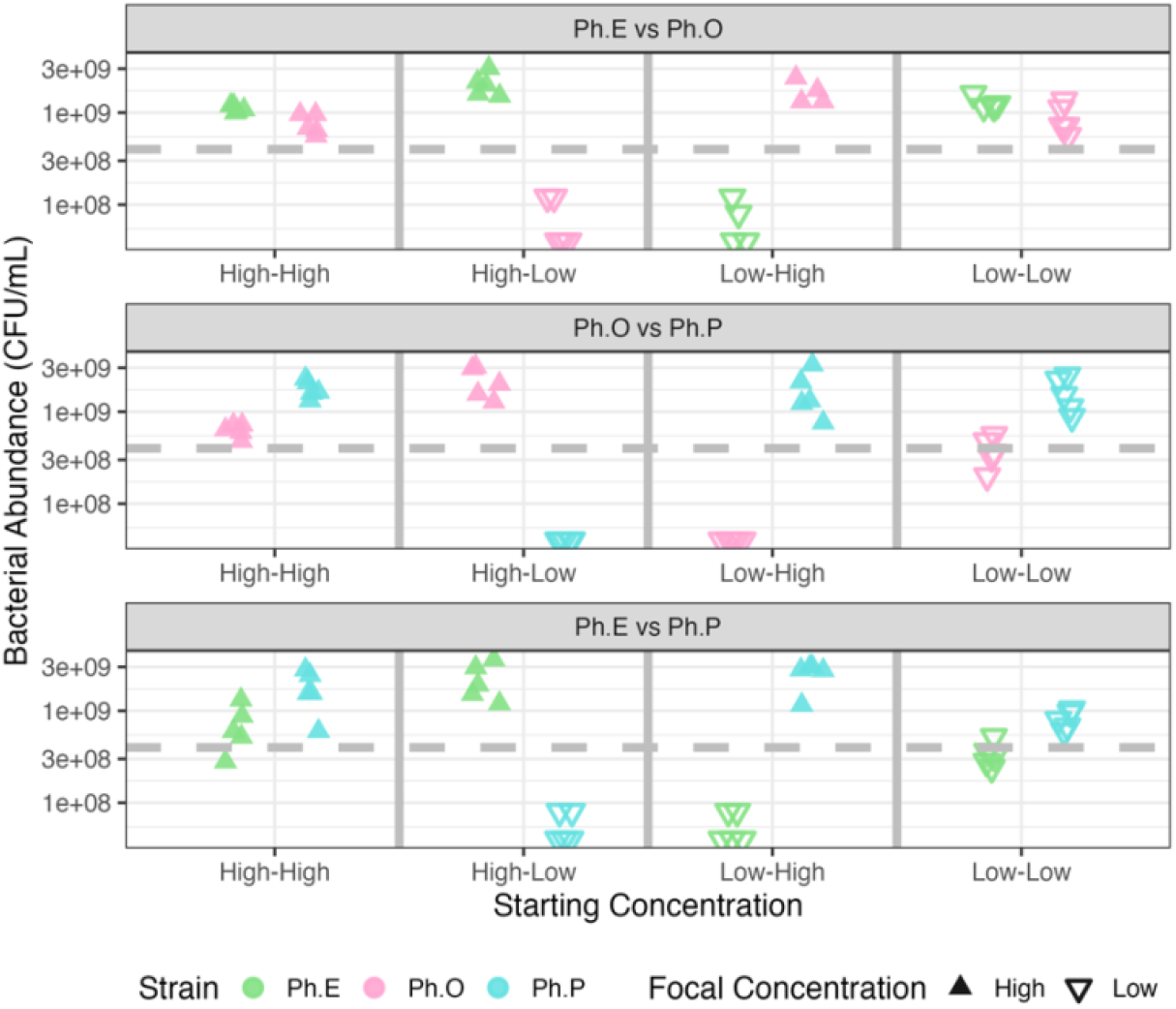
Abundance of *Photobacterium mandapamensis* strains Ph.E (green), Ph.O (pink), and Ph.P (blue) when grown in pairwise communities after 10 hours under different initial conditions. Strain abundance was measured by colony counts. The dashed line indicates the threshold of 10 colonies, below which counts were considered unreliable for calculating CFU/mL. Plates with zero colonies were assigned a value of 1 to reflect very low abundance rather than true absence. The treatment conditions are indicated on the X axis. Upward-pointing triangles denote strains that were introduced at a higher concentration; downward-pointing triangles indicate strains at a lower concentration.

**Figure S3:**
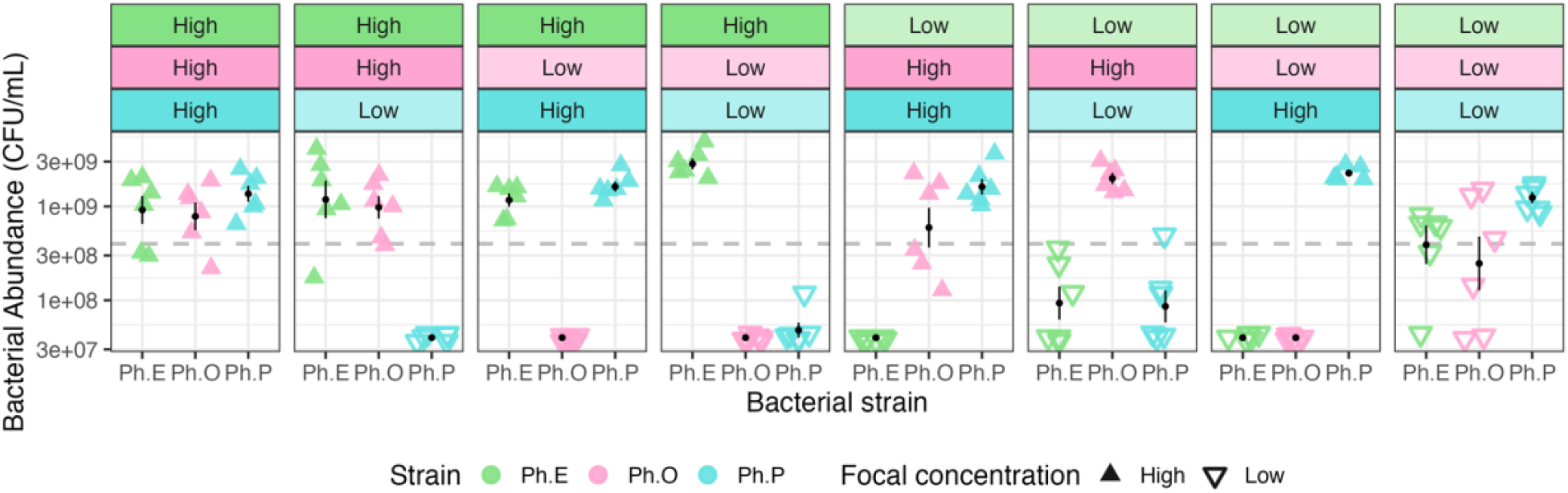
Abundance of *Photobacterium mandapamensis* strains Ph.E (green), Ph.O (pink), and Ph.P (blue) when grown in mixed communities after 10 hours under different initial conditions. Strain abundance was measured by colony counts. The dashed line indicates the threshold of 10 colonies, below which counts were considered unreliable for calculating CFU/mL. Plates with zero colonies were assigned a value of 1 to reflect very low abundance rather than true absence. The treatment conditions are indicated above each panel. Upward-pointing triangles denote strains that were introduced at a higher concentration; downward-pointing triangles indicate strains at a lower concentration.

**Figure S4:**
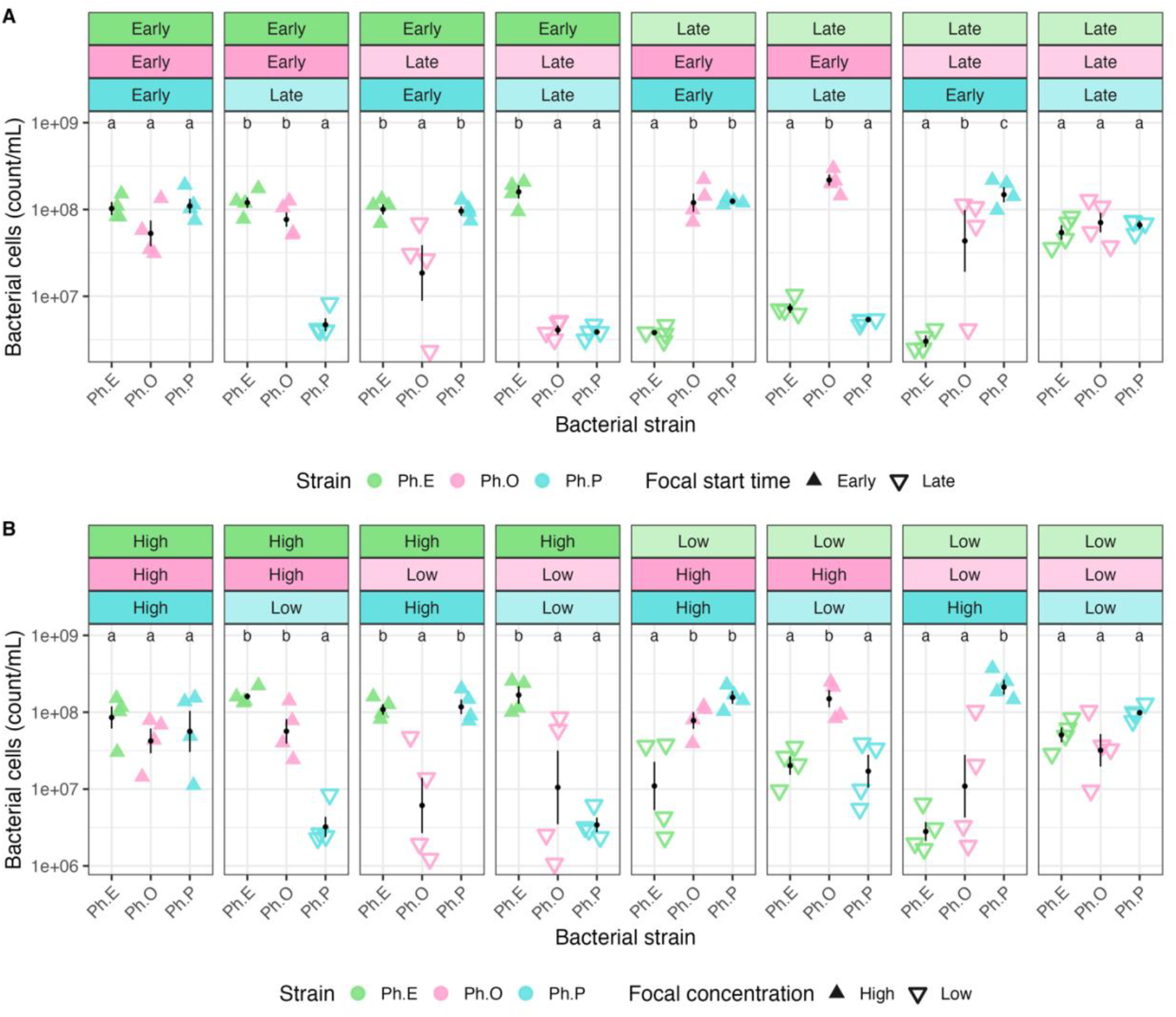
Abundance of *Photobacterium mandapamensis* strains Ph.E (green), Ph.O (pink), and Ph.P (blue) when grown in mixed communities after 24 hours under different initial conditions. Strain abundance was measured using flow cytometry. Priority effects were simulated by staggering arrival order (A) or altering initial strain concentrations as a proxy for arrival order (B). The treatment conditions are indicated above each panel. Upward-pointing triangles denote strains that arrived first or were introduced at a higher concentration; downward-pointing triangles indicate strains that arrived later or at a lower concentration. Each point represents the average abundance across biological replicates (n = 4), with black points representing means ± standard error indicated by black bars. Strains sharing the same letter are not significantly different from one another. In both experimental setups, strains introduced first or at higher concentrations maintained higher abundance relative to their competitors. These results show priority effects have a strong role in shaping strain-level community composition.

## Notes

### Competing Interest Statement

The authors have declared no competing interest.

### Summary of Updates

Name of authors revised and spelling updated.

## Sources

1. Bai B et al. The root microbiome: Community assembly and its contributions to plant fitness. Journal of Integrative Plant Biology 2022;64:230–243. 10.1111/jipb.13226

2. Costello EK et al. The application of ecological theory toward an understanding of the human microbiome. Science 2012;336:1255–1262. 10.1126/science.1224203

3. Coyte KZ et al. Ecological rules for the assembly of microbiome communities. PLOS Biology 2021;19:e3001116. 10.1371/journal.pbio.3001116

4. Gould AL et al. Microbiome interactions shape host fitness. Proceedings of the National Academy of Sciences 2018;115:E11951–E11960. 10.1073/pnas.1809349115

5. Fukami T. Historical Contingency in Community Assembly: Integrating Niches, Species Pools, and Priority Effects. Annual Review of Ecology, Evolution, and Systematics 2015;46:1–23. 10.1146/annurev-ecolsys-110411-160340

6. Boyle JA et al. Priority effects alter interaction outcomes in a legume–rhizobium mutualism. Proceedings of the Royal Society B: Biological Sciences 2021;288:20202753. 10.1098/rspb.2020.2753

7. Chappell CR et al. Wide-ranging consequences of priority effects governed by an overarching factor. eLife 2022;11:e79647. 10.7554/eLife.79647

8. Cheong JZA et al. Priority effects dictate community structure and alter virulence of fungal-bacterial biofilms. ISME J 2021;15:2012–2027. 10.1038/s41396-021-00901-5

9. Debray R et al. Priority effects in microbiome assembly. Nat Rev Microbiol 2022;20:109–121. 10.1038/s41579-021-00604-w

10. Devevey G et al. First arrived takes all: inhibitory priority effects dominate competition between co-infecting Borrelia burgdorferi strains. BMC Microbiology 2015;15:61. 10.1186/s12866-015-0381-0

11. Jones KR, Belden LK, Hughey MC. Priority effects alter microbiome composition and increase abundance of probiotic taxa in treefrog tadpoles. Applied and Environmental Microbiology 2024;90:e00619–24. 10.1128/aem.00619-24

12. Laursen MF, Roager HM. Human milk oligosaccharides modify the strength of priority effects in the Bifidobacterium community assembly during infancy. The ISME Journal 2023;17:2452–2457. 10.1038/s41396-023-01525-7

13. Ojima MN et al. Priority effects shape the structure of infant-type Bifidobacterium communities on human milk oligosaccharides. The ISME Journal 2022;16:2265–2279. 10.1038/s41396-022-01270-3

14. Pucci N et al. Priority effects, nutrition and milk glycan-metabolic potential drive Bifidobacterium longum subspecies dynamics in the infant gut microbiome. PeerJ 2025;13:e18602. 10.7717/peerj.18602

15. Shao Y et al. Primary succession of Bifidobacteria drives pathogen resistance in neonatal microbiota assembly. Nat Microbiol 2024;9:2570–2582. 10.1038/s41564-024-01804-9

16. Gould AL, Harii S, Dunlap PV. Host preference, site fidelity, and homing behavior of the symbiotically luminous cardinalfish, Siphamia tubifer (Perciformes: Apogonidae). Mar Biol 2014;161:2897–2907. 10.1007/s00227-014-2554-z

17. Gould AL et al. Life history of the symbiotically luminous cardinalfish Siphamia tubifer (Perciformes: Apogonidae). Journal of Fish Biology 2016;89:1359–1377. 10.1111/jfb.13063

18. Dunlap PV et al. Symbiosis initiation in the bacterially luminous sea urchin cardinalfish Siphamia versicolor. Journal of Fish Biology 2012;81:1340–1356. 10.1111/j.1095-8649.2012.03415.x

19. Dunlap PV, Nakamura M. Functional morphology of the luminescence system of Siphamia versicolor (Perciformes: Apogonidae), a bacterially luminous coral reef fish. Journal of Morphology 2011;272:897–909. 10.1002/jmor.10956

20. Gould AL et al. Strain-level diversity of symbiont communities between individuals and populations of a bioluminescent fish. ISME J 2023;17:2362–2369. 10.1038/s41396-023-01550-6

21. Urbanczyk H et al. Genome Sequence of Photobacterium mandapamensis Strain svers.1.1, the Bioluminescent Symbiont of the Cardinal Fish Siphamia versicolor ?. J Bacteriol 2011;193:3144–3145. 10.1128/JB.00370-11

22. Gould A, Henderson J. Highly contiguous genome assemblies of Photobacterium strains isolated from fish light organs using nanopore sequencing technology. 2022. 2022.

23. Bongrand C et al. Using Colonization Assays and Comparative Genomics To Discover Symbiosis Behaviors and Factors in Vibrio fischeri. mBio 2020;11:10.1128/mbio.03407-19. 10.1128/mbio.03407-19

24. Bongrand C, Ruby EG. Achieving a multi-strain symbiosis: strain behavior and infection dynamics. The ISME Journal 2019;13:698–706. 10.1038/s41396-018-0305-8

25. Bongrand C, Ruby EG. The impact of Vibrio fischeri strain variation on host colonization. Current Opinion in Microbiology 2019;50:15–19. 10.1016/j.mib.2019.09.002

26. Galardini M et al. Phenotype inference in an Escherichia coli strain panel. eLife 2017;6:e31035. 10.7554/eLife.31035

27. Lloyd-Price J et al. Strains, functions and dynamics in the expanded Human Microbiome Project. Nature 2017;550:61–66. 10.1038/nature23889

28. Yan Y et al. Strain-level epidemiology of microbial communities and the human microbiome. Genome Med 2020;12:71. 10.1186/s13073-020-00765-y

29. Gould AL, Osland HK. Strain-level variation in microbial symbiosis: lessons from the Siphamia–Photobacterium mandapamensis system. Trends in Microbiology 2025;0. 10.1016/j.tim.2025.02.010

30. Alker AT et al. A modular plasmid toolkit applied in marine bacteria reveals functional insights during bacteria-stimulated metamorphosis. mBio 2023;14:e01502–23. 10.1128/mbio.01502-23

31. Alker AT et al. Linking bacterial tetrabromopyrrole biosynthesis to coral metamorphosis. ISME Communications 2023;3:98. 10.1038/s43705-023-00309-6

32. Sprouffske K, Wagner A. Growthcurver: an R package for obtaining interpretable metrics from microbial growth curves. BMC Bioinformatics 2016;17:172. 10.1186/s12859-016-1016-7

33. Noh S et al. Facultative symbiont virulence determines horizontal transmission rate without host specificity in Dictyostelium discoideum social amoebas. Evol Lett 2024;8:437–447. 10.1093/evlett/qrae001

34. Peay KG, Belisle M, Fukami T. Phylogenetic relatedness predicts priority effects in nectar yeast communities. Proc R Soc B 2012;279:749–758. 10.1098/rspb.2011.1230

35. Garud NR et al. Evolutionary dynamics of bacteria in the gut microbiome within and across hosts. PLoS Biol 2019;17:e3000102. 10.1371/journal.pbio.3000102

36. Verster AJ et al. The Landscape of Type VI Secretion across Human Gut Microbiomes Reveals Its Role in Community Composition. Cell Host & Microbe 2017;22:411-419.e4. 10.1016/j.chom.2017.08.010

37. Wollenberg MS, Ruby EG. Population Structure of Vibrio fischeri within the Light Organs of Euprymna scolopes Squid from Two Oahu (Hawaii) Populations. Applied and Environmental Microbiology 2009;75:193–202. 10.1128/AEM.01792-08

38. Chen JZ et al. A strong priority effect in the assembly of a specialized insect-microbe symbiosis. Applied and Environmental Microbiology 2024;90:e00818–24. 10.1128/aem.00818-24

39. Wang S et al. Chemolithoautotrophic diazotrophs dominate dark nitrogen fixation in mangrove sediments. The ISME Journal 2024;18:wrae119. 10.1093/ismejo/wrae119

40. Dunlap PV, Urbanczyk H. Luminous Bacteria. In: Rosenberg E et al. (eds.), The Prokaryotes: Prokaryotic Physiology and Biochemistry. Berlin, Heidelberg: Springer, 2013, 495–528.

41. Widder EA. Bioluminescence in the Ocean: Origins of Biological, Chemical, and Ecological Diversity. Science 2010;328:704–708. 10.1126/science.1174269

42. Louca S et al. Function and functional redundancy in microbial systems. Nat Ecol Evol 2018;2:936–943. 10.1038/s41559-018-0519-1

43. Wolff R, Shoemaker W, Garud N. Ecological Stability Emerges at the Level of Strains in the Human Gut Microbiome. mBio 2023;14:e02502–22. 10.1128/mbio.02502-22

44. Nyholm SV, McFall-Ngai MJ. A lasting symbiosis: how the Hawaiian bobtail squid finds and keeps its bioluminescent bacterial partner. Nat Rev Microbiol 2021;19:666–679. 10.1038/s41579-021-00567-y

45. Sun Y et al. Intraspecific Competition Impacts Vibrio fischeri Strain Diversity during Initial Colonization of the Squid Light Organ. Appl Environ Microbiol 2016;82:3082– 3091. 10.1128/AEM.04143-15

